# Sox5 controls the establishment of quiescence in neural stem cells during postnatal development

**DOI:** 10.1101/2024.05.03.592315

**Authors:** Cristina Medina-Menéndez, Lingling Li, Paula Tirado-Melendro, Pilar Rodríguez-Martín, Elena Melgarejo-de la Peña, Mario Díaz-García, María Valdés-Bescós, Rafael López-Sansegundo, Aixa V. Morales

## Abstract

Adult stem cells niches relays in the acquisition of a reversible state of quiescence to ensure long-lasting DNA integrity and cell expansion. Neural stem cells (NSCs) in the dentate gyrus (DG) enter quiescence before the adult hippocampal neurogenic niche is fully established. However, the mechanisms controlling NSC first quiescence entry and quiescence deepness are largely unknown. Using conditional mutant mouse during embryonic or postnatal stages, we have determined that transcription factor Sox5 is required to restrict first entry in quiescence. Moreover, we have found a critical window during the second postnatal week when NSCs build up a shallow or primed quiescent state. Loss of Sox5 leads to an excess of primed NSCs prone to activate leading to a neurogenic burst in the adult DG and precocious depletion of the NSC pool. Mechanistically, Sox5 prevent an excess of BMP canonical signaling activation, a pathway that we have now determined is associated to NSC primed state. In conclusion, our results demonstrate that Sox5 is required to control the correct balance between primed and deep quiescence during the first postnatal weeks of DG development, a balance which is essential for establishing long-lasting adult neurogenesis.

## Introduction

During embryonic and postnatal development of the dentate gyrus (DG), neural stem cells (NSCs) proliferate, migrate and generate mature granule neurons (GN) and astrocytes. In sharp contrast to other brain regions, in the subgranular zone (SGZ) of the adult DG a subpopulation of NSCs remains in a reversible quiescent state (qNSCs) [1–4]. Through highly regulated molecular mechanisms, those adult NSCs are activated (aNSCs) and enter in cell cycle to produce intermediate progenitor cells (IPCs) that generate new GNs in the DG throughout adult life [5,6].

NSCs in the DG enter in a quiescence state predominantly during the first postnatal week and a high percentage of them will remain as a source of adult new neurons [1,3]. Recently, it has been shown that autophagy drives the conversion of developmental NSCs to the adult quiescent state [7]. However, little is known about the transcriptional program controlling this first quiescence entry and how the reversibility of the initial quiescent state is achieved.

Furthermore, quiescence is not an unique static state defined by cell cycle absence. Single-cell RNA-Seq analysis of adult NSCs support the idea of a continuum of cell states or NSC populations from a deep/dormant quiescent state to an activated state [8–10]. This includes a shallow, resting or primed quiescent state defined, in comparison with the dormant state, by higher activation rate, upregulation of ribosomal genes and a shift in energy metabolism genes [8–10]. Moreover, the primed state can be captured in culture where combinations of cell cycle markers define primed NSCs (pNSCs) at a half way position between qNSCs and aNSCs [11,12]. However, it is unclear how NSCs acquire this primed state as they enter quiescence and convert into adult NSCs during the first postnatal weeks.

Despite the complexity of signalling pathways acting within the NSC adult niche, candidates for promoting adult NSC quiescence have emerged. These include BMP4 [13,14] and its downstream effectors Id1 and Id4 [15–17]; Delta/Notch pathway [18–20] and the MFGE8/integrin/ILK pathway [21], amongst others. Nevertheless, we know very little about signals and transcription factors that could modulate quiescence during DG development.

Now, using mouse conditional mutant for Sox5 during embryonic or early postnatal development we describe a critical window around P14 when NSCs build up a shallow or primed quiescent state. During that period, Sox5–defective mice exhibit an increase in pNSCs that lead to severe alterations in the adult DG neurogenic niche, including transient aberrant NSC activation and excessive neurogenesis. As a consequence, older *Sox5* mutant mice show a premature reduction in the adult NSC pool and in neurogenesis. Mechanistically, Sox5 prevent an excess of BMP/P-Smad1/5/9/Id4 activation, a canonical pathway, which we found, is associated to the primed state in NSCs.

## Results

### NSC population and proliferation decrease drastically during the second postnatal week of DG development

During the first postnatal week the majority of NSCs in the developing DG enter quiescence for their first time [1,3] and most adult GNs are generated [22,23]. In order to systematically quantitate possible changes in the NSC population during postnatal development, we identified NSCs by GFAP and Sox2 expression and determined NSC proliferating fraction using MCM2 (a cell cycle marker) from P0 to P150 (Fig. 1A-D). We observed a dramatic NSC loss during the first postnatal week (P0>P5) from 4.16×10^6^ to 1.1×10^6^ cells/mm^2^ (Fig. 1A, B) accounting for a depletion rate of 19.9 % of NSCs per day (Fig. 1C). However, we observed that NSC proliferation rate (% of MCM2^+^ NSCs) remained stable during that period (Fig. 1D). The reduction in the NSC pool during P0>P5 interval could be due to the peak in GN differentiation previously described [22,23] and supported by the increase in Prox1^+^ GNs over total DG cell population from 56.5 ± 2.5% to 75.8 ± 3% (Fig. 1E).

**Fig. 1.**
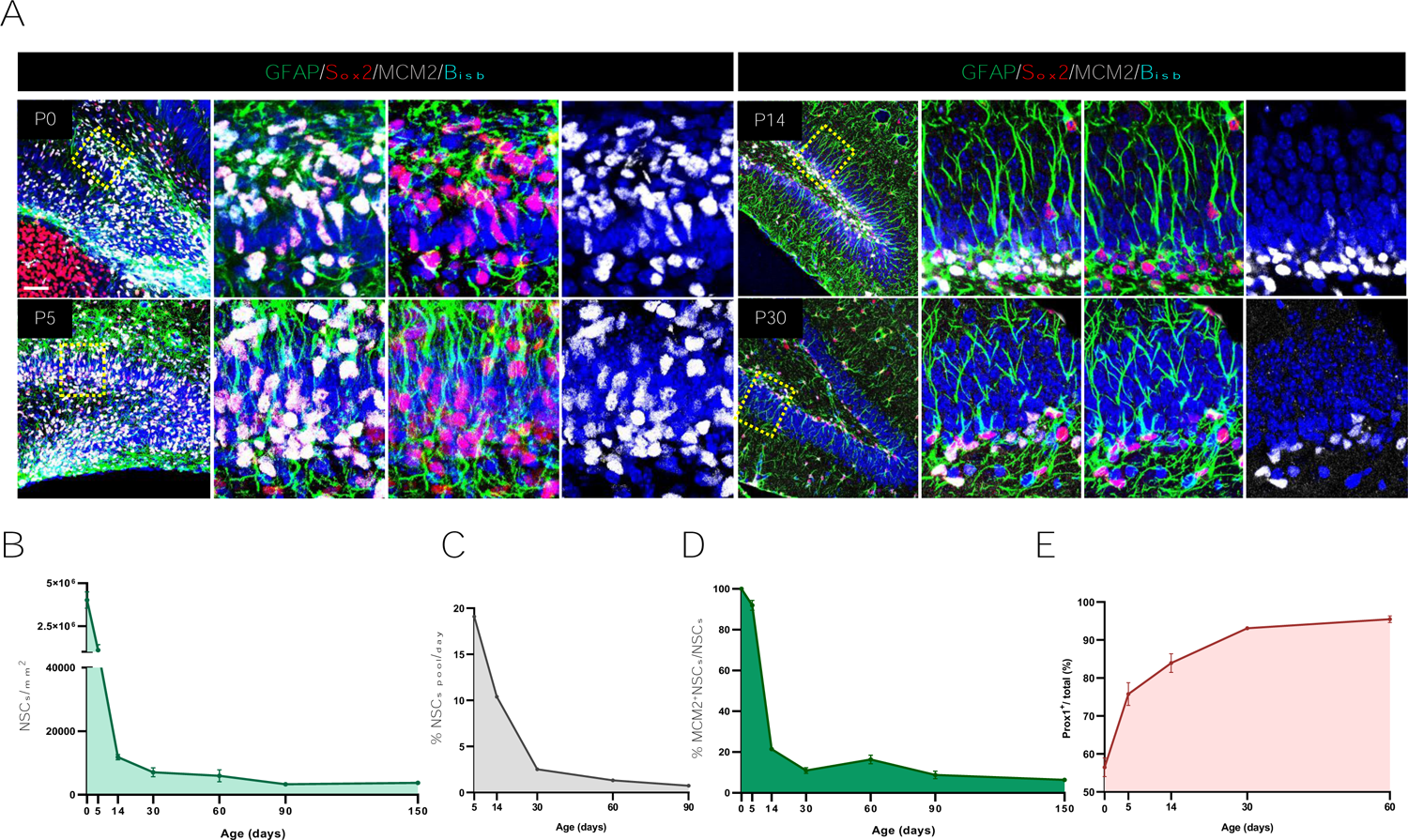
NSC population and proliferation decrease drastically during the second postnatal week of DG development. **(A)** Confocal images showing proliferative NSCs (rGFAP^+^Sox2^+^MCM2^+^) in the DG at the indicated mice stages. **(B)** Quantitation of NSCs/mm^2^ in the DG (rGFAP^+^Sox2^+^ cells) at different stages. **(C)** NSC depletion rate per day in DG. **(D)** Proliferating NSC quantitation in the NSC population (%). **(E)** Quantitation of Prox1^+^cells amongst total DG cells (%). At least 3 animals and 3 sections/animal were analyzed for each immunostaining. In all graphs, data are mean value ± SEM. ***p < 0.001 by unpaired Student’s t test. Scale bars represent 30 µm (A, right panels), 100 µm (A left panels)

By the second postnatal week, NSC loss continued (from 1.1×10^6^ to 11846.5 cells/mm^2^ at P5>P14; Fig. 1A, B), albeit at a slightly lower depletion rate (10.4% daily loss; Fig. 1C). Nevertheless, during this second week we observed a dramatic loss of the NSC proliferating fraction (only 21.5 ± 0.9% of the NSC pool remained by P14), whereas Prox1^+^ cells increased to 83.9 ± 2.5%. These data suggest that by the end of the second postnatal week the depletion of the NSC pool could be due both to NSC differentiation and to NSC accumulation in a quiescent state that prevents new rounds of NSC self-renewal.

At early adulthood, the pool of NSCs declined sharply by P30 at a rate of 2.6% per day and at a moderate rate by P60 and P90 (Fig. 1A-C). In parallel, the proliferating fraction halved (10.9 ± 1.4% of MCM2^+^ NSCs; Fig. 1A, D) with respect to that in P14 and then remained similar by P60 and P90 (16.4 ± 2.1% and 8.8 ± 1.9%; Fig. 1D). This coincides with the majority of DG cells becoming Prox1^+^ GNs by P30 (93. 1± 0.3%) and P60 (95.5 ± 0.9%; Fig. 1E).

Thus, there are two temporal windows of high NSC variation: i) P0>P5, when the NSC pool is severely reduced mostly due to massive neuronal differentiation and ii) P5>P14, when the NSC pool decreases possibly due both to cell differentiation and to massive NSC accumulation in quiescence, hindering NSC self-renewal.

### Sox5 is required to restrict NSC entry in quiescence during the first postnatal week

In order to understand early dynamics in NSC population, we analysed the expression of transcription factor Sox5 that is required for the transition from quiescence to activation in the adult DG [24]. We observed that Sox5 expressing cells were widely distributed in the developing hilus and granular zone (GZ) of the DG from embryonic day 16.5 to P5 (Fig. 2C and Fig. S1). At P5, NSCs are accumulated in the future SGZ and the majority of Sox5^+^ cells in the DG co-expressed NSC markers Sox2, GFAP and Hopx and were proliferating (Fig. 2C and Fig. S1C, D). Moreover, we estimated that 9.6 ± 0.8 % of Sox2^+^ cells were Tbr2^+^ IPCs and almost none of them were Prox1^+^ GNs (Fig. S1C, E). Considering the high level of Sox5 and Sox2 co-expression, we could assume that Sox5 was expressed in around 10% of IPCs and almost absent from GNs.

**Fig 2.**
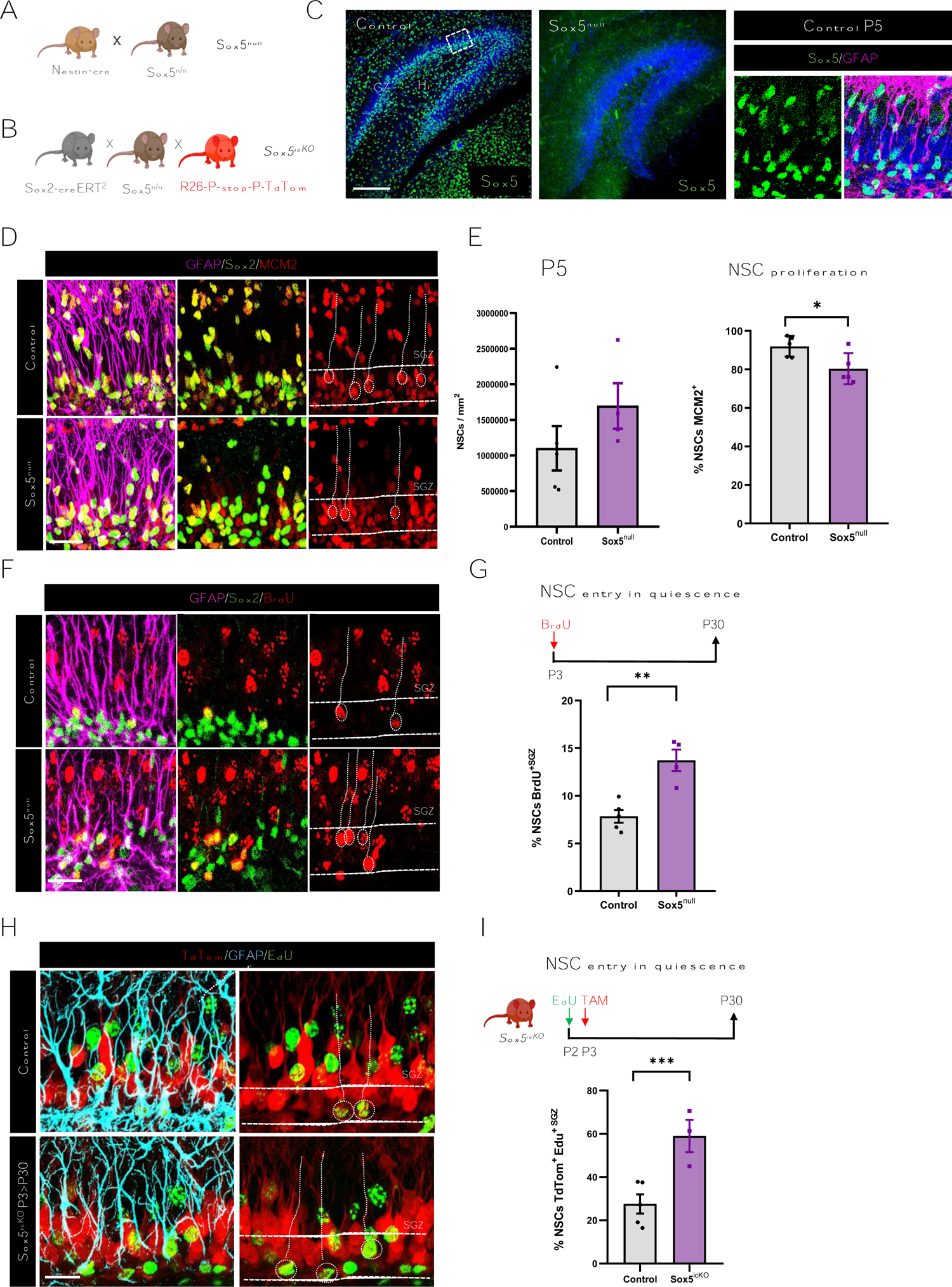
Sox5 is required to restrict NSC entry in quiescence during the first postnatal week. **(A, B)** Generation of Sox5^null^ (A) and Sox5^icKO^ (B) mice. **(C)** Confocal images showing Sox5 expression in dorsal DG of Control and Sox5^null^ mice by P5 and Sox5^+^ cells expressing GFAP in Control P5 DG. **(D)** Confocal images showing GFAP, Sox2 and MCM2 immunostaining in the SGZ (space between white lines) of P5 Control and Sox5^null^ mice. **(E)** Quantitation of NSCs/mm^2^ (rGFAP^+^Sox2^+^ cells) and proliferative MCM2^+^ NSCs (rGFAP^+^Sox2^+^MCM2^+^ % over total NSCs) in the SGZ of Control and Sox5^null^ P5 mice. **(F)** Confocal images showing GFAP, Sox2 and BrdU immunostaining in P30 Control and Sox5^null^ mice after P3 BrdU pulse. **(G)** Scheme of BrdU-long labelling retention experiment. Quantitation of BrdU^+^ NSCs % in the BrdU^+^ population of P30 Control and Sox5^null^ mice SGZ. **(H)** Confocal images showing TdTom, GFAP and EdU immunostaining in the SGZ of P30 Control and Sox5^icKO^ mice. **(I)** Scheme of EdU-long labeling retention experiment in TAM injected conditional Sox5^icKO^ mice (P3>P30). Quantitation of EdU^+^ NSCs % in the EdU^+^TdTom^+^ population of P30 Control and Sox5^icKO^ mice SGZ. Data represent mean values ± SEM. *p < 0.05, **p < 0.01, and ***p < 0.001 by unpaired Student’s t test. Scale bars represent 100 µm (C) and 50 µm (D, F, H)

By P14, the majority of Sox5^+^ cells were radial (r) GFAP^+^/Sox2^+^ cells but more than half of them stopped proliferating (only 36.1 ± 6.1% were MCM2^+^) and only 57.2 ± 3.5% retained Hopx (Fig. S1C, D). Hopx expression is progressively restricted to quiescent NSCs during postnatal DG development [25]. Thus, Sox5^+^Hopx^+^ cells could represent the increasing pool of quiescent NSCs by P14. Further analysis showed that 8.2 ± 0.9% of Sox2^+^ cells were Tbr2^+^ IPCs and 3.5 ± 1.7% were Prox1^+^ GNs (Fig. S1C, E). In conclusion, Sox5 expressing cells in the postnatal developing DG were predominantly NSCs and Sox5 expression decreases as NSCs progress along the neurogenic cascade.

To determine if Sox5 was required for NSC entry into quiescence for their first time, we resourced to a conditional Sox5^fl/fl^ line [26] crossed to a Nestin-cre line to remove Sox5 expression from neural progenitors starting around embryonic stage 13.5 (Fig. 2A, C; Sox5^null^). During the first postnatal week, we observed that loss of Sox5 reduced NSC proliferation with respect to Control mice (80.4 ± 3.6% vs. 92.0± 2.4% of MCM2^+^ cells in rGFP^+^/Sox2^+^ NSCs; *P* = 0.029; Fig. 2D, E). However, the NSC pool was similar in Sox5^null^ and Control mice (Fig. 2E).

To explore if the reduction in aNSCs in Sox5^null^ mice was due to an abnormal entry in quiescence, we performed a single BrdU injection at P3 and chased NSCs by P30 (Fig. 2G). Dividing NSCs can incorporate BrdU and retain it for as long as they remain quiescent [BrdU-long (BrdU-L) retaining NSCs]. By P30, the percentage of BrdU-L NSCs with respect to total BrdU^+^ cells was larger in Sox5^null^ than in Control mice (13.7 ± 1.1% vs. 7.9 ± 0.7% respectively; *P* = 0.0023; Fig. 2F, G). Thus, embryonic Sox5 loss provoked postnatal increase in the number of NSCs entering quiescence and a reduction in aNSCs.

As neural progenitors of the embryonic DG express Sox5 (Fig. S1A), Sox5^null^ NSC defects at P5 could be due to embryonic alterations. To clarify this aspect, we resourced to a conditional inducible strategy using a Sox2-creER^T2^ mouse line crossed with Sox5^fl/fl^ and reporter Rosa-TdTomato lines previously characterized (Sox5^icKO^; Fig. 2B) [24]. Sox5 loss induced by TAM injection at P3, preceded by a single EdU injection, caused a dramatic increase in recombinant TdTom^+^ EdU-Long retaining NSCs by P30 with respect to Control mice (59.0 ± 7.5% vs 27.6 ± 4.5% respectively, *P* = 0.0082; Fig. 2H, I). As these results were similar to those observed in Sox5^null^ mice (Fig. 2F, G), we concluded that Sox5 is required during the first postnatal week to prevent an excess of NSCs entering quiescence for their first time and thus to maintain NSCs in a proliferative state which ensures adequate NSC number.

### Embryonic loss of Sox5 reduces the pool of dormant quiescent NSCs by P14

To determine if Sox5 was required during the second postnatal week to prevent an excess of NSCs entering quiescence, we promoted selective Sox5 loss at P14 by TAM injection in Sox5^icKO^ mice. Similarly, to what we had described in P3>P30, by P14>P30 Sox5^icKO^ mice exhibited a robust increase in the number of recombined TdTom^+^ EdU ^+^ NSCs with respect to Control mice (91.40 ± 4.8% vs. 59.7 ± 9.5% respectively, *P* = 0.0118; Fig. 3F, G), indicating an increase in NSCs entering quiescence by P14 due to postnatal Sox5 loss.

**Fig. 3.**
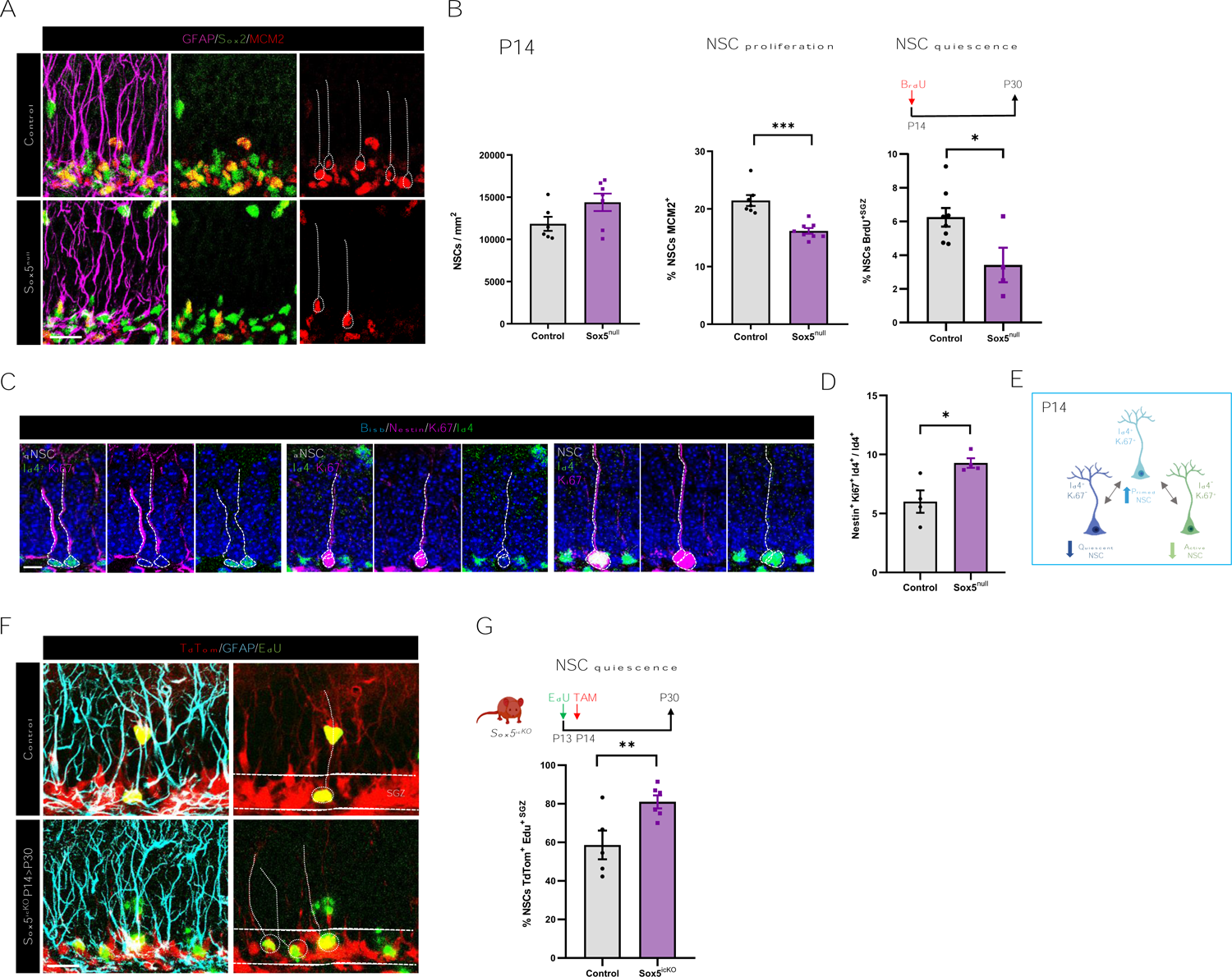
Embryonic loss of Sox5 reduces the pool of quiescent NSCs by P14. **(A)** Confocal images showing GFAP, Sox2 and MCM2 immunostaining in the SGZ (space between white lines) of P14 Control and Sox5^null^ mice. **(B)** Quantitation of NSCs/mm^2^ (rGFAP^+^Sox2^+^ cells), proliferative MCM2^+^ NSCs (% over total NSCs) in Control and Sox5^null^ P14 mice. Scheme of BrdU-long labelling retention experiment. Quantitation of BrdU-L^+^ NSCs % in the BrdU^+^ population of P30 Control and Sox5^null^ mice SGZ. **(C)** Confocal images showing Nestin, Ki67 and Id4 immunostaining in the SGZ of P14 Control and Sox5^null^ mice. **(D)** Quantitation of NSCs Nestin^+^ Id4^+^Ki67^+^ over total Id4^+^ cells in Control and Sox5^null^ P14 mice. **(E)** Schematic summary of the alterations in qNSC, pNSC and aNSC balance in P14 Sox5^null^ mice. **(F)** Confocal images showing TdTom, GFAP and EdU immunostaining in the SGZ of P30 Control and Sox5^icKO^ mice. **(G)** Scheme of EdU-long labeling retention experiment in TAM injected conditional Sox5^icKO^ mice (P14>P30). Quantitation of EdU^+^ NSCs % in the EdU^+^TdTom^+^ population of P30 Control and Sox5^icKO^ mice SGZ. In all graphs, data represent mean value ± SEM. *p < 0.05, **p < 0.01, and ***p < 0.001 by unpaired Student’s t test. Scale bars represent 50 µm (A, C, G) and 10 µm (E)

Conversely, by P14 Sox5^null^ mice exhibited a reduction in aNSCs % with respect to Control mice (16.2 ± 0.5% vs. 21.5 ± 1.0% of MCM2^+^ cells in the population of rGFAP^+^ Sox2^+^; *P* = 0.0002; Fig. 3A, B) and the NSC pool did not changed (Fig. 3B).These results indicate that Sox5 is required for NSC proliferation during the first two weeks of postnatal DG development.

Unexpectedly, in Sox5^null^ mice injected with BrdU at P14 and chased by P30 we observed a reduction in BrdU-L NSCs in comparison with Control mice (3.4 ± 1.0% vs. 6.3 ± 0.6% of BrdU^+^ rGFAP^+^ Sox2^+^ in BrdU^+^ pool; *P* = 0.0226; Fig. 3B). Thus, although the immediate response of TAM-induced-Sox5 loss is the increase in quiescence entry (Fig. 3G), the mid-term response to Sox5 loss in Sox5^null^ mice is a reduction in NSCs that remain in quiescence from P14 to P30. However, if both aNSC and qNSC populations are reduced in Sox5^null^ mice (Fig. 3B) and the NSC pool remains unchanged, in which state are the remaining NSCs?

To approach the puzzle, we analysed the expression of the marker of quiescence Id4 [13,14] in P14 Nestin^+^ NSCs. As expected, in Control mice the majority of Id4^+^ cells were Ki67^-^Nestin^+^ qNSCs whereas aNSCs could be identified as Nestin^+^Ki67^+^Id4^-^ cells. However, we observed few Nestin^+^ NSCs that were double positive (Ki67^+^Id4^+^) and that could represent an intermediate state between quiescence and activation, previously defined as resting or primed state [10]. Surprisingly, Sox5^null^ mice showed an increase in these Nestin^+^ Ki67^+^ Id4^+^ NSCs in comparison with Control mice (9.3 ± 0.4 vs. 6.0 ± 0.9 %; *P* = 0.0196; Fig. 3C, D).Thus, these data suggest that around P14, there is a subpopulation of Id4^+^ NSCs, probably in an intermediate state between activation and quiescence, which is selectively increased by Sox5 loss (Fig. 3E).

### Sox5 loss causes a general over activation of NSCs in young adult mice

If there were an excess of NSCs in an intermediate or primed state at P14 in Sox5^null^ mice, it could potentially compromise the normal activation/quiescence balance in the young adult neurogenic niche. In fact, in P30 Sox5^null^ mice we observed that the % of aNSCs was surprisingly higher than in Control mice (18.8 ± 1.4 vs. 10.9 ± 1.4% of MCM2^+^cells in the pool of rGFAP^+^ Sox2^+^ cells; *P* = 0.0018; Fig. 4A, B). Moreover, using Sox5^icko^ mouse to delete Sox5 expression postnatally we confirmed that P30 NSCs showed a higher rate of proliferation in Sox5^icko^ than in Control mice from P3>P30 (16.3 ± 1.4% vs. 10.1 ± 0.9%; *P*= 0.0056; Fig. 4D) or from P14>P30 (24.9 ± 1.3% vs. 14.2 ± 1.7 %; *P* = 0.0015; Fig. 4C, D). These data demonstrate that loss of Sox5, either during development or at different postnatal weeks, leads to the same output in the established young adult DG neurogenic niche: an excess of young adult aNSCs at P30.

**Fig. 4.**
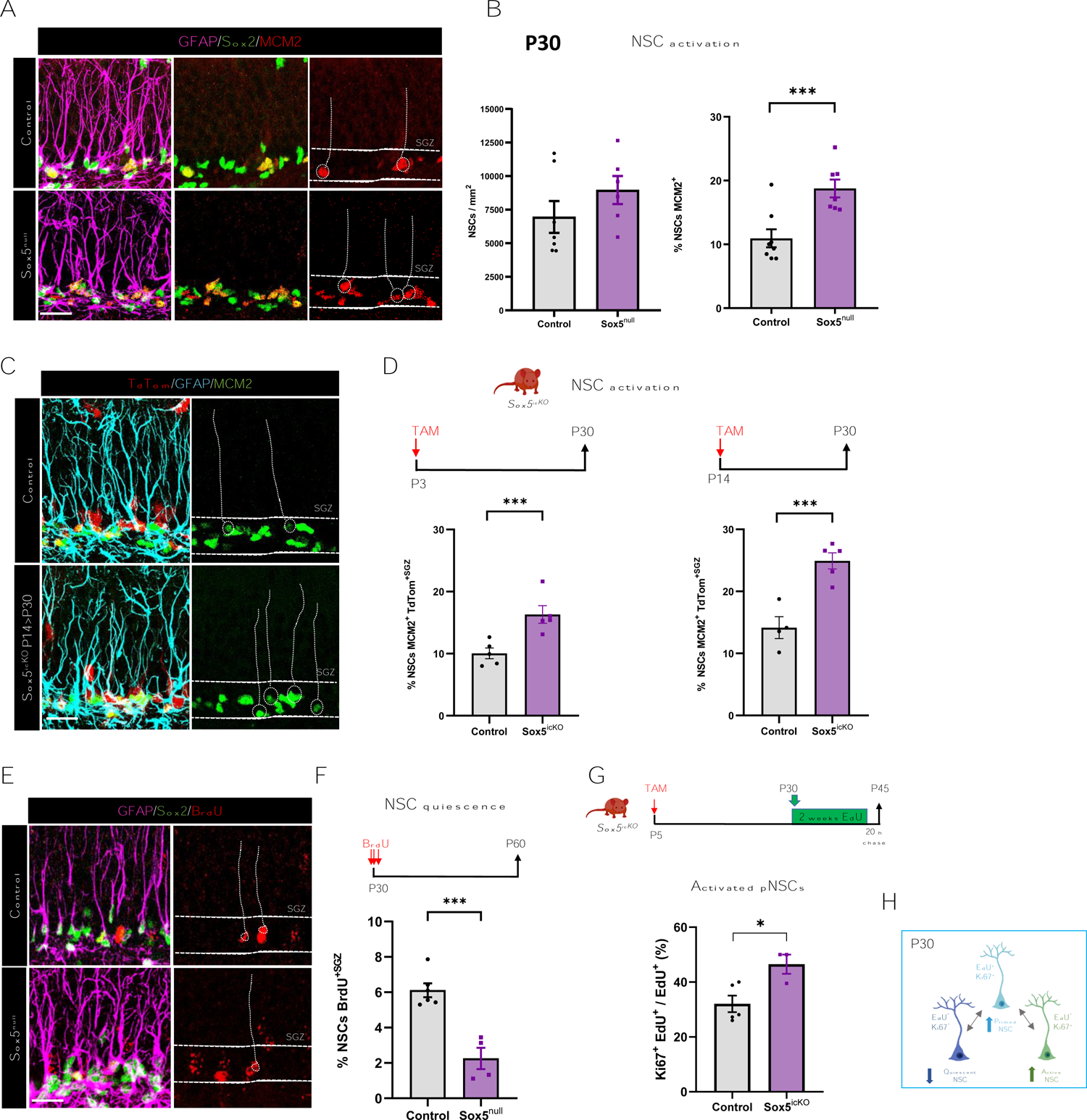
Sox5 loss causes an over activation and reduced return to quiescence of NSCs in young adult mice. **(A)** Confocal images showing GFAP, Sox2 and MCM2 immunostaining in the SGZ (space between white lines) of P30 Control and Sox5^null^ mice. **(B)** Quantitation of NSCs/mm^2^ (rGFAP^+^Sox2^+^cells) and proliferative MCM2^+^ NSCs (% over total NSCs) in the SGZ of Control and Sox5^null^ P30 mice. **(C)** Confocal images showing TdTom, GFAP and MCM2 immunostaining in the SGZ of P30 Control and Sox5^icKO^ mice. **(D)** Scheme of Sox5 inducible deletion at P3>P30 and at P14>P30 in Sox5^icKO^ mice by TAM injection. Quantitation of aNSCs (rGFAP^+^MCM2^+^TdTom^+^) in the TdTom^+^ population in SGZ of P30 Control and Sox5^icKO^ mice. **(E)** Confocal images showing GFAP, Sox2 and BrdU immunostaining in the SGZ of P60 Control and Sox5^null^ mice. **(F)** Scheme of BrdU-long label retention experiment. Quantitation of BrdU-L^+^ NSCs % in the BrdU^+^ population of P30 Control and Sox5^null^ mice SGZ. **(G)** Experimental scheme of EdU label retention experiment in P5>P30 in Sox5^icKO^ mice to label all pNSCs (EdU^+^) during P30>P45 period. EdU^-^ NSCs correspond to qNSCs. Quantitation of % pNSCs that have been activated (EdU^+^ Ki67^+^) in the total EdU^+^Nestin^+^TdTom^+^ population of P45 Control and Sox5^icKO^ mice. In all graphs, data represent mean values ± SEM. *p < 0.05, **p < 0.01, and ***p < 0.001 by unpaired Student’s t test. Scale bars represent 50 µm (A, C and E). **(H)** Schematic summary of the alterations in qNSC, pNSC and aNSC balance in P30 Sox5^null^ mice.

Conversely, using BrdU-L retention assays in Sox5^null^ mice from P30 to P60 we observed a clear decrease in the proportion of BrdU-L^+^ NSCs with respect to Control mice (2.3 ± 0.6% vs 6.1 ± 0.4%, respectively; *P* = 0.0005; Fig. 4E, F) indicating a lower return to quiescence of the aNSCs in Sox5^null^ mice. These results reinforce the data that young adult NSCs in Sox5^null^ P30 mice are more engaged in activation and less in remaining in quiescence than Control NSCs.

To determine if this reduction in qNSCs from P30 to P60 was accompanied by changes in NSCs in a primed state, we resourced to a saturating EdU retention assay [10]. EdU was administered in drinking water at P30 for two weeks followed by 20-hour chase before culling at P45 in combination with Ki67 immunolabelling (Fig. 4G). We could identify pNSCs as those that have incorporated and retained EdU pulse and the dormant quiescent pool as those NSCs that remained EdU^-^ during the two-week period. Upon Sox5 loss by TAM injection at P5 we observed in P45 Sox5^icKO^ mice an increase in activated pNSCs (46.5 ± 3.5 vs. 32.1 ± 2.8 % of EdU^+^Ki67^+^ NSCs in the EdU^+^TdTom^+^ Nestin^+^ NSC pool; *P* = 0.028; Fig. 4G) in comparison with Control mice. In other words, pNSCs showed a lower tendency to return to quiescence with respect to those in Control mice (Fig. S2). In summary, our results suggest that loss of Sox5 provokes, in the young adult neurogenic niche, an increase in pNSCs with impaired return to quiescence after activation and the over-activation of NSCs (Fig. 4H).

### Over activation of NSCs provokes a transient increase in neurogenesis in young adult Sox5^null^ mice and a premature depletion of the NSC pool

In order to determine if the excess in activation of young adult NSCs in Sox5^null^ mice could have any consequence in neurogenesis, we analysed the number of DCX^+^ immature neurons. At P14 and P23, a decrease in immature neurons was detected in Sox5^null^ with respect to Control mice (3214 ± 124 vs 4295 ± 289 DCX^+^ BrdU^+^ cells/mm^2^; *P* = 0.04; Fig. S3 and 1.3 ×10^6^ ± 7.5 ×10^4^ vs. 1.5 ×10^6^ ± 7.0 ×10^4^ cells/mm^3^; *P* = 0.05; Fig. 5B). However, by P30 new neurons doubled in Sox5^null^ mice with respect to control mice (1.2 ×10^6^ ± 0.1 ×10^6^ vs. 0.6 ×10^6^ ± 0.1 ×10^6^ cells/mm^3^, respectively; *P* = 0.0009; Fig. 5A, B), indicating that by P30 the excess of aNSC described (Fig. 4B) could be responsible for an increase in neurogenesis in Sox5^null^ mice.

**Fig. 5.**
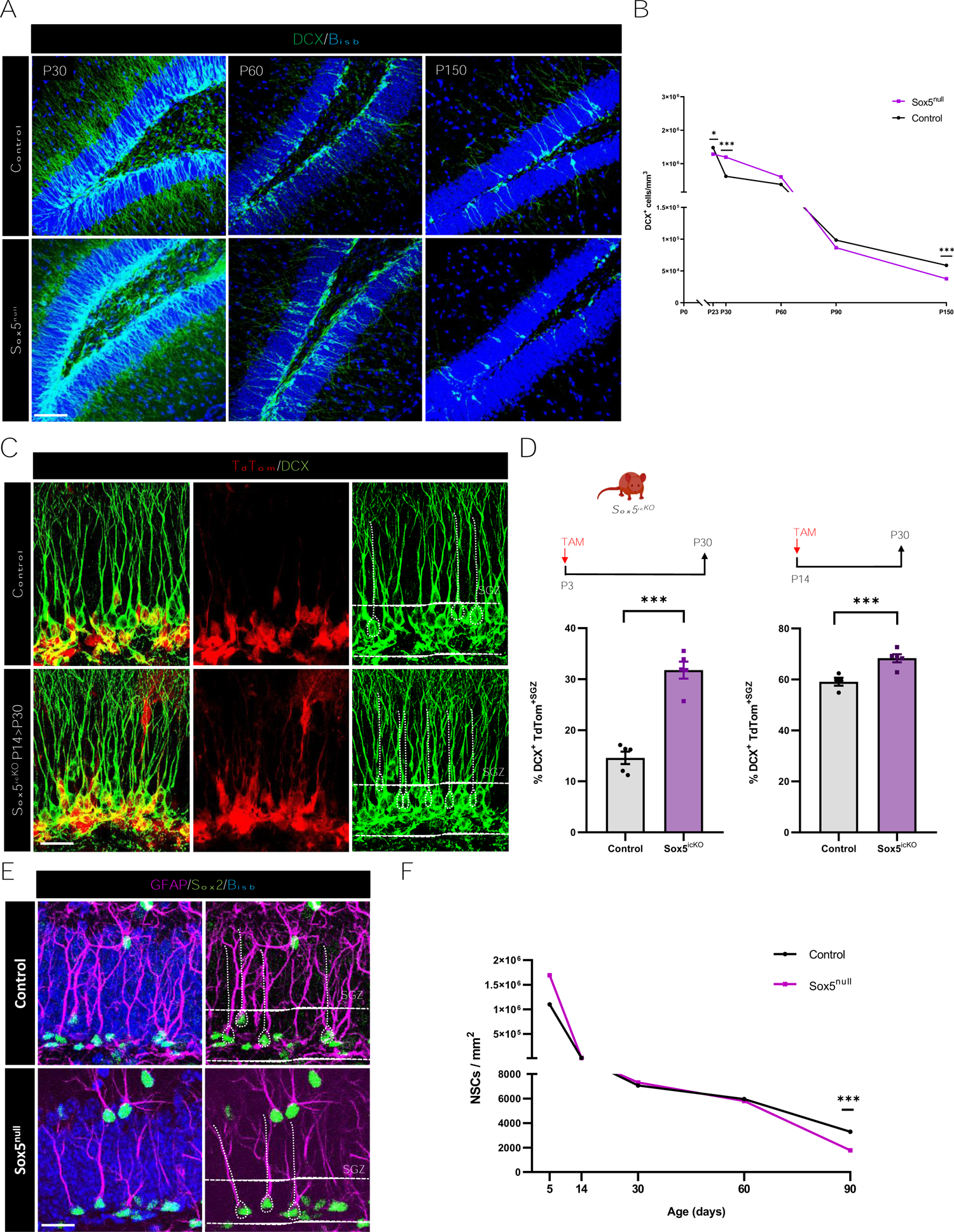
Overactivation of NSCs provokes a transient increase in neurogenesis in young adult Sox5^null^ mice and a premature depletion of the NSC pool. (A) Confocal images showing DCX immunostaining in the SGZ of DG at the indicated stages in Control and Sox5^null^ mice. (B) Time-course quantitation of DCX^+^ cells/mm^3^ at the indicated stages in Control and Sox5^null^ mice. (C) Confocal images showing TdTom^+^ and DCX^+^ immunostaining in the SGZ of P30 Control and Sox5^icKO^ mice. (D) Scheme of Sox5 inducible deletion at P3>P30 and at P14>P30 in Sox5^icKO^ mice by TAM injection. Quantitation of % TdTom^+^DCX^+^ cells over total TdTom^+^ cells in the SGZ in P30 Control and Sox5^icKO^ mice. (E) Confocal images showing GFAP and Sox2 immunostaining in the SGZ of P90 Control and Sox5^null^ mice. (F) Quantitation of NSCs (rGFAP^+^Sox^2+^ cells)/mm^2^ at the indicated stages in Control and Sox5^null^ mice. At least 3 animals and 3 sections/animal were analysed for each immunostaining. In all graphs, data are mean value ± SEM. *p < 0.05, **p < 0.01, and ***p < 0.001 by unpaired Student’s t test. Scale bars represent 100 µm (A) and 30 µm (C, E)

To confirm that this defect was not due to embryonic changes in neuronal cell fate decisions in Sox5^null^ mice, we promoted postnatal Sox5 loss at P3 and P14 by TAM-induced recombination in Sox5^icko^. Again, we observed a robust increase in DCX^+^ neurons in the TdTom^+^ recombined cell population both at P3>P30 and P14>P30 Sox5^icko^ mice in relation with Control mice (31.8± 1.7 % vs. 14.6 ± 1.2 %, *P* <0.0001 and 68.3 ± 1.6% vs. 59.1 ± 1.6%, *P* = 0.005, respectively; Fig. 5C, D). Thus, loss of Sox5 before the establishment of the adult neurogenic niche (either at embryonic or early postnatal stages) leads to an excess in neuronal production in young adult mice by P30.

Next, we explored if the excess of adult neurogenesis in P30 remained along Sox5^null^ mice life. We did not observe significant changes in the number of DCX^+^ cells at P60 and P90 between Sox5^null^ and Control mice (Fig. 5A, B), indicating that the excess of neurogenesis at P30 could be compensated later on. Unexpectedly, by P150 the number of new neurons significantly decreased in Sox5^null^ with respect to Control mice (37618 ± 6786 vs. 59956 ± 5948 cells/mm^3^, *P* = 0.04; Fig. 5A, B), indicating that Sox5 is required for long-term neurogenesis.

There are certain situations such as loss of pro-quiescence genes (Mfge8) [21], epileptogenic conditions [27] or BMP inhibition in ageing mice [28], in which NSC over-activation and excess in neuronal production eventually leads to NSC depletion. In fact, although there were not changes in the size of the NSC pool from P14 to P60 between Control and Sox5^null^ mice, by P90 we detected a dramatic depletion of the adult NSC pool in mutant mice with respect to Control (1780 ± 242 vs. 3299 ± 180 cells/mm^2^, *P* =0.0013; Fig. 5E, F). In summary, these results indicate that NSC over-activation in young P30 Sox5^null^ mice, leads to a temporal excess in neuronal production that later derives in a premature depletion of the NSC pool and diminished neurogenesis. Moreover, our results suggests that early control of the right balance of activated, primed and quiescent states is crucial for the long-term maintenance of the adult neurogenic niche.

### Transcriptomic analysis of Sox5^null^ DG reveals dramatic changes in developmental programs in the adult neurogenic niche, including TGF-β/BMP pathway activation

To elucidate the molecular mechanisms behind the profound alterations in NSC activation and neuron production observed upon Sox5 loss, we performed RNA sequencing (RNA-Seq) comparative analysis in Control and Sox5^null^ DG at stage P34 (n= 4 and 3, respectively; Fig. 6A). We first established by principal component analysis (PCA) of differentially expressed genes (DEGs; adjusted p-value < 0.05; Deseq^2^ software) that our individual samples segregated clearly into Control and Sox5^null^ groups (Fig. 6B). Furthermore, the heat map of DEGs clearly showed individual differences in the level of expression of 2961 genes between groups (Fig. 6C). Similarly, the distribution in a volcano plot showed 1572 genes upregulated (Fold Change, FC> +0.5) and 1389 genes downregulated (FC< −0.5) in the DG of Sox5^null^ mice with respect to Control ones (Fig. 6D). To capture biological information on DEG, Gene Ontology (GO) analysis for biological processes highlighted changes in development (system development, developmental processes, anatomical structure morphogenesis or nervous system development; Fig. 6E). These DEGs could be at the molecular base of the changes in NSC proliferation and aberrant neurogenesis previously described (Figs. 4 and 5). Similarly, GO analysis for cellular components highlighted changes in neuronal specific components such as synapse, neuron projection, dendritic tree, axon or presynapse components that closely matched the phenotypic analyses of Sox5^null^ DG (Fig. 6E).

**Fig. 6.**
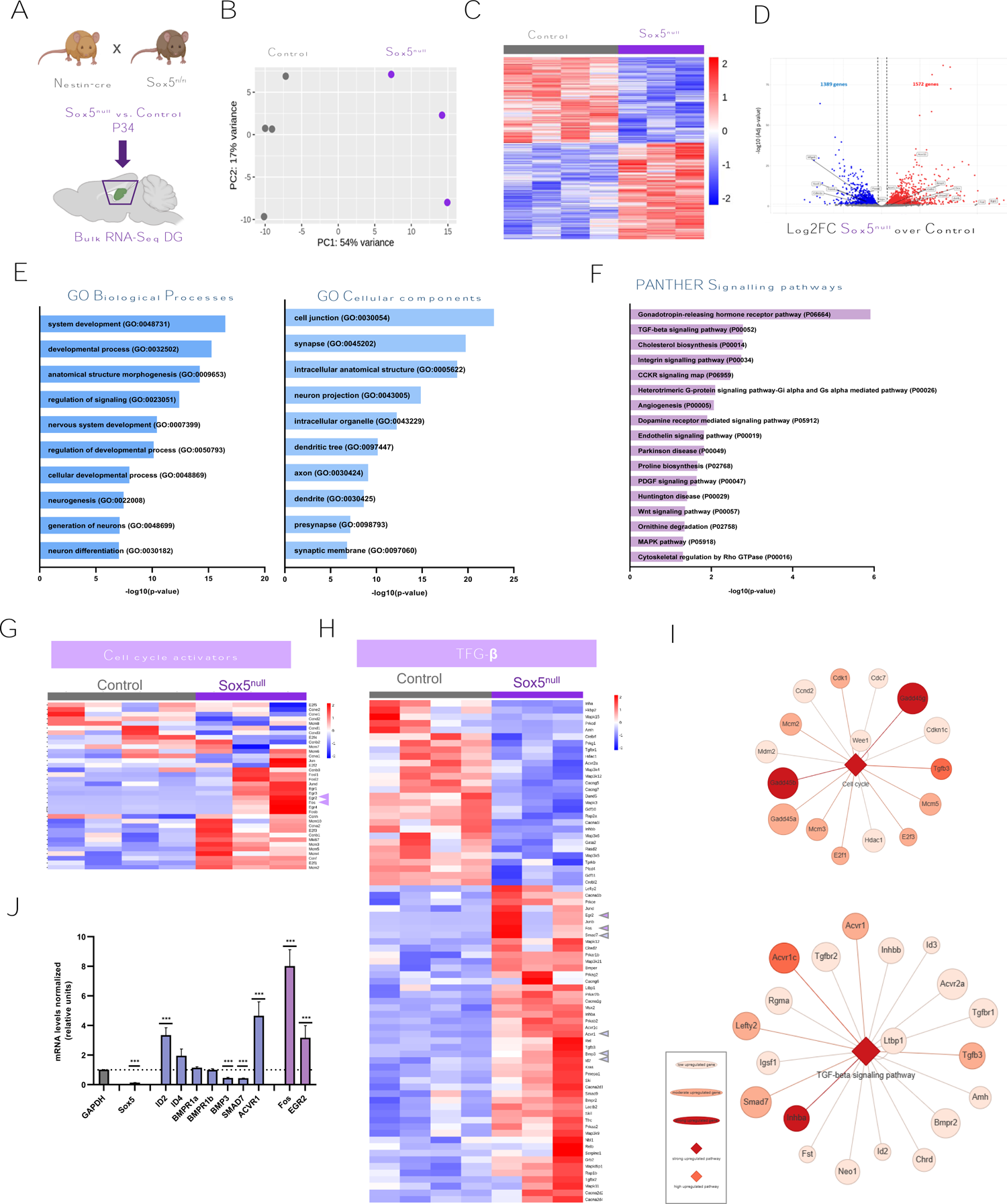
Transcriptomic analysis of Sox5^null^ DG reveals important changes in developmental programs in the adult neurogenic niche including activation of TGF-β superfamily signalling pathway. (A) Experimental approach of bulk RNA-Seq analysis of P34 DG. (B) Principal Component Analysis of DEGs indicating independent segregation of the two genotypes analyzed, Control (grey) and Sox5^null^ (violet). (C) Heat Map of DEGs in Sox5^null^ vs Control. (D) Volcano Plot of LogFC and –log10 of all tested DEGs between Control and Sox5^null^ mice. Several relevant genes are indicated in boxes and those related to quiescence (*Mfge8, Hopx, Foxo6*) are downregulated whereas those related to neurogenesis (*Dcx, Calb2, Egr2, Fos, Junb*) are upregulated in Sox5^null^ vs Control DG. (E) Enrichment analysis of GO terms related to Biological Processes and Cellular Components in the DGEs of Sox5^null^ vs Control DG. (F) Enrichment analysis of GO terms related to signalling pathways in DGEs of Sox5^null^ vs Control DG. DGEs were selected for their p-value < 0.05 and the Fisher’s Exact type test was used together with Bonferroni’s multiple testing correction by the open access software Panther (v.14.0). (G-H) Heat Map of DEGs manually curated in lists of cell cycle activators (G) or TFG-β/BMP signalling pathway genes (H) comparing Sox5^null^ vs Control DG. (I) Node graphs of most representative upregulated cell cycle activators and TGF-β signalling pathway genes generated by the PANEV software (PAthway NEtwork Visualizer; v. 17.0). (J) Quantitation of the relative levels of mRNA expression by quantitative PCR for the indicated transcripts in Sox5^null^ vs Control DG. Results are shown as 2^-ΔΔCT^ normalized with respect to *GAPDH* mRNA and relative to P30 Control values (dashed line y-axis=1). Four independent samples were analyzed for each condition and experimental replicates were performed. Data represent mean values ± SEM. ***p<0.001 according to a Student’s t-test

Using Panther Software to explore possible alterations in signalling pathways upon Sox5 loss, we found a cluster of TGF-β signalling as one of the top ranked. This pathway has been strongly associated to NSC quiescence regulation [14,29] and to neuronal differentiation [30] (Fig. 6F). We also found signalling pathways associated with NSCs proliferation such as Wnt and MAPK pathways [31,32] (Fig. 6F). In order to determine if the combination of DEGs were indicative of up or downregulation of each of the pathways they were associated to, we resourced to PANEV (PAthway NEtwork Visualizer) [33] a recently described R package set for gene/pathway-based network visualization based on information available on KEGG. We tested several of the Panther identified signalling pathways and observed a high upregulation of cell cycle genes, especially *Mcm* genes such as *Mcm2*, *Mcm3* and *Mcm5* and the TGF-β pathway (Fig. 6I).

These results were also supported by heat map representation of 36 preselected genes described in the bibliography as cell cycle activators including *Cyclins*, *Mcms*, *Fos/Jun* and *Egrs* genes (Fig. 6G) observing that the majority of the activators were upregulated. These data were further confirmed by qPCR, showing an increase in Sox5^null^ P34 DGs of *Fos* and *Egr2* expression with respect to Control mice (8.0 ± 1.1-folds, *P*= 0.003 and 3.2 ± 0.8, *P*= 0.003, respectively; Fig. 6J). Similarly, we could also observe that in a cluster of 76 TGF-β signalling related genes the majority showed a clear upregulation trend in Sox5^null^ DGs (Fig. 6H) that was confirmed by qPCR for TGF-β/BMP targets such as *Id2* (3.4 ± 0.5-folds, *P* = 0.009) and the activin receptor *Acvr1* (4.5 ± 0.9-folds *P*= 0.017). In addition, we confirmed that, as in the RNA-Seq, there were not significant changes by qPCR in the expression of *Id4* (1.9 ± 0.5 -folds, *P* = 0.2), *Bmpr1a* (1.1 ± 0.1-folds, *P* = 0.1) and *Bmpr1b* (1.0 ± 0.08-folds, *P*= 0.9) in Sox5^null^ mice in comparison with Control mice (Fig. 6J). In summary, bulk transcriptomic analyses revealed that Sox5 loss of expression during DG development causes marked changes in gene expression at the onset of adult neurogenesis, providing a comprehensive view of the excess in cell cycle activation and neuronal differentiation in the neurogenic niche. Moreover, these studies point to the relevance TGF-β/BMP pathway has in the control of early NSC quiescence.

### Sox5 is required to prevent an excess of Id4^+^ primed NSCs during the second postnatal week

To further explore if possible changes in pNSCs and in TGF-β or BMP pathways could be behind those alterations observed in P14 Sox5^null^ mice, we resourced to an *in vitro* model in which hippocampus-derived NSCs were grown as floating neurospheres in the presence of mitogens (Fig. 7A) [24]. To capture activated and quiescent states, P14 hippocampal NSCs expanded by several passages were loaded with a cell tracer (DFFDA-Oregon Green fluorophore) that are diluted over cell divisions (Fig. 7A) [11]. In this culture, dormant quiescent NSCs neither engage in cell cycle nor survive past 48 hour [11,34], whereas cells in a primed state retain high levels of DFDDA and contribute predominantly to the long-term maintenance of the NSC culture [11]. After six days in culture in the presence of FGF2, by flow cytometry analysis, we determined that the majority of P14 NSCs diluted the cell-tracer upon cell division (79.8± 7.2 %, DFFDA^low^) whereas few cells retained high DFDDA levels (19.6 ± 7.2 %, DFFDA^high^; Fig. 7B, C). Moreover, adding BMP4 that promotes quiescence entry [14] in combination with FGF2, we observed a distinct shift in the cell profile as 80.4 ± 5.3 % of cells were DFFDA^high^ and only 19.3 ± 5.3 % were DFFDA^low^ (Fig. 7B, C). We then compared Sox5^null^ and Control P14 NSCs and observed a significant increase in the % of DFFA^high^ cells in Sox5^null^ NSCs grown in FGF2 compared to Control (20.0 ± 2.5% vs 9.9 ± 0.4%; *P* = 0.007; Fig. 7F). These results indicate that Sox5 loss promotes the generation of a higher number of primed-like NSCs identified as DFFDA^high^ cells.

**Fig. 7.**
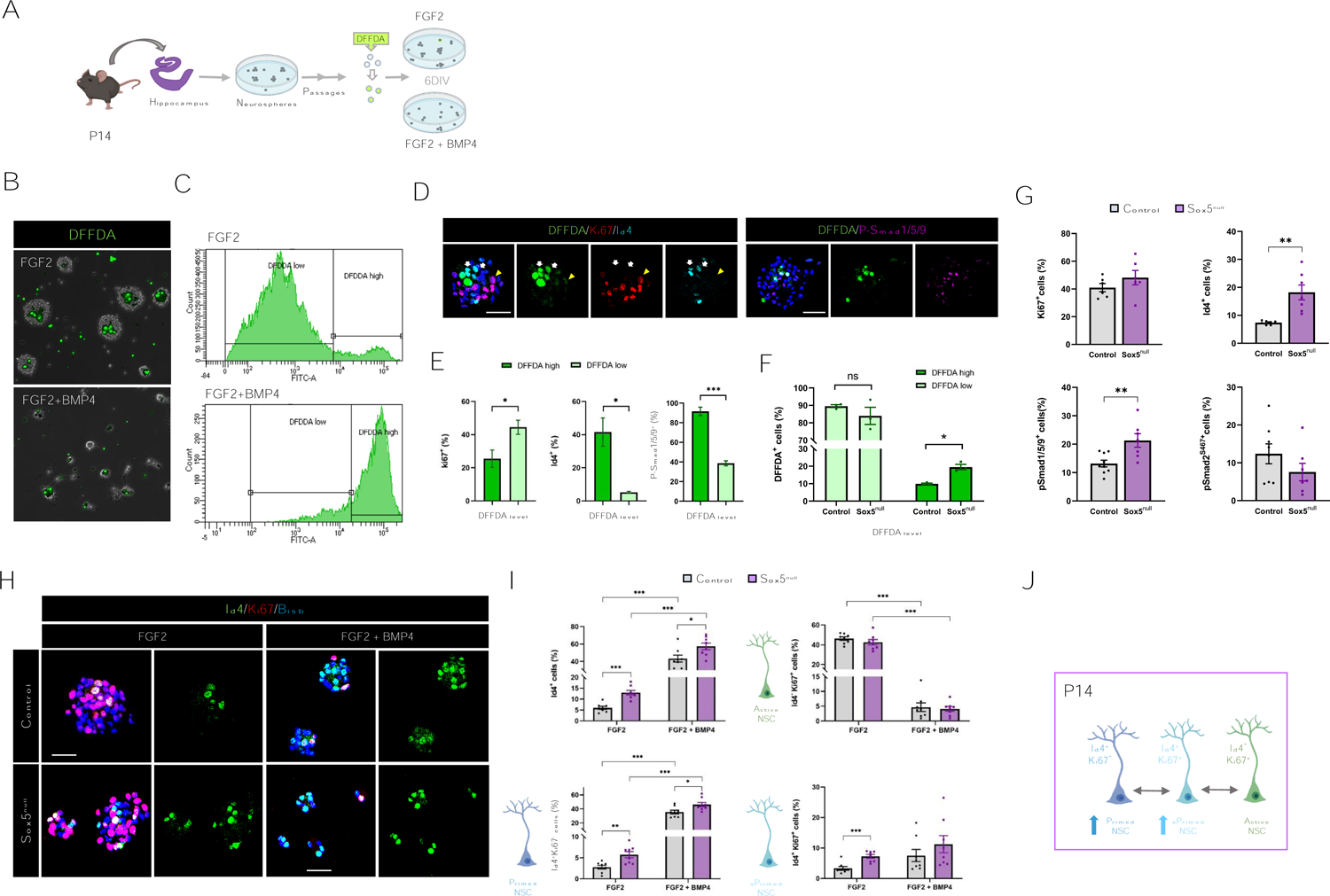
Sox5 is required to prevent an excess of primed quiescent Id4^+^ NSCs during the second postnatal week. **(A)** Methodological approach for NSC culture. **(B)** Representative bright field images of P14 neurospheres loaded with DFFDA (green) under proliferating (FGF2) and quiescence (FGF2+BMP4) culture conditions for 6 days. **(C)** Quantitation using flow cytometry of DFFDA^high^ and DFFDA^low^ cells at the indicated culture condition. **(D)** Confocal images showing DFFDA fluorescence and Ki67, Id4 or P-Smad1/5/9 expression in P14 NSCs. **(E)** Quantitation of the percentage of P-Smad1/5/9^+^, Ki67^+^ and Id4^+^ cells among DFFDA^high^ and DFFDA^low^ cells from Control mice cultured in FGF2. **(F)** Quantitation of DFFDA^high^ and DFFDA^low^ cells in Control and Sox5^null^ mice in proliferation. **(G)** Quantitation of the percentage of P-Smad1/5/9^+^, Ki67^+^ and Id4^+^ cells relative to total NSC number in Control and Sox5^null^ mice cultured in FGF2 (passages 2-6).**(H)** Confocal images showing Ki67 and Id4 immunostaining in the indicated conditions. **(I)** Quantitation of the percentage of Id4^+^ cells, Id4^+^/Ki67^-^ (pNSCs), Id4^-^/Ki67^+^ (aNSCs) and Id4^+^/Ki67^+^ (activated pNSCs) relative to total cell number in Control and Sox5^null^ mice. **(J)** Schematic summary of the alterations in pNSC and aNSC balance in P14 Sox5^null^ mice DG *in vitro*.Data represent mean values ± SEM. * p<0.05; ** p<0.01; ***p<0.001 according to a Student’s t-test. Scale bars represent 100 µm (B) and 30 µm (G, E)

As BMP canonical pathway has been associated to the quiescent state [14,35] and we had observed that Id4^+^/Ki67^+^ cells could define the primed NSC population *in vivo* (Fig. 3C-E), we tested several BMP canonical targets that could be associated with the primed state of NSCs *in vitro*. Thus, we found that Id4 (a BMP signalling effector) was enriched in P14 DFFDA^high^ cells with respect to DFFDA^low^ cells (41.5 ± 8.6 % vs 5.2 ± 0.5 %; *P*= 0.05; Fig. 7D, E). That was also the case for phosphorylated Smad1/5/9 (P-Smad1/5/9, a mediator and direct indicator of BMP canonical signalling activation; 91.5 ± 4.3 % vs 38.6 ± 2.5 %; *P*= 0.0005; Fig. 7D, E), whereas the opposite distribution was observed for cell cycle marker Ki67 (25.4 ± 5.2% vs 44.5 ± 4.2 %; *P*= 0.047; Fig. 7D, E). These results indicate that pNSCs (DFFDA^high^) *in vitro* are characterized by high levels of BMP canonical pathway activation.

In Sox5^null^ DG we had observed an upregulation of components of BMP canonical pathway (e.g. *Bmpr2* and *Id2*), TGF-β canonical pathway (e.g.*Tgfbr2* and *Tgfbr1*) and Activin canonical pathway (*Acvr1* and *Acvr2a;* Fig. 6H-J). In order to determine which of those pathways could be involved in the excess of pNSCs observed in Sox5^null^ mice (Figs. 3C-E and 7F), we used acute NSCs preparation from P14 DG, from passage 0 (primary) to passage 6. We did not observed significant changes in the number, size and proliferation (Ki67^+^ cells) of Sox5^null^ NSCs with respect to those of Control mice (Fig. S5 and Fig. 7G). However, we observed a consistent upregulation of BMP canonical pathway mediator P-Smad1/5/9 and effector Id4 in Sox5^null^ NSCs with respect to Control NSCs (21.3 ± 2.4 % vs 13.2 ± 1.2 %; *P*= 0.006 and 18.2 ± 2.6 % vs 7.4 ± 0.3 %; *P*= 0.006, respectively; Fig. 7G), whereas TGF-β/Activin canonical pathway mediator P-Smad2 was not significantly affected (Fig. 7G). Thus, Sox5 is required to supress BMP canonical pathway in postnatal NSCs.

After expanding control and mutant NSCs for several passages (5 to 15 passages) to obtain enough NSC number, we explored Sox5^null^ NSCs responsiveness to a canonical BMP pathway ligand such as BMP4. In Control NSCs and as expected for a BMP target, the number of Id4^+^ cells dramatically increased when NSCs were exposed to BMP4+FGF2 with respect to FGF2 condition (43.3 ± 3.9 % vs 6.0 ± 0.7 %; *P* < 0.0001; Fig. 7H, I). Interestingly, NSCs derived from Sox5^null^ P14 mice showed a higher proportion of Id4^+^ NSCs with respect to Control mice both in FGF2 (13.0 ± 1.0 % vs 6.0 ± 0.7 %; *P* < 0.0001; Fig. 7H, I) and BMP4+FGF2 conditions (57.3 ± 3.8 % vs. 43.3 ± 3.9 %; *P* = 0.02; Fig. 7H, I), reinforcing the idea that Sox5 prevents an excess of BMP canonical activation both in proliferating and quiescence conditions.

Moreover, when classifying NSCs according to Ki67 and Id4 expression, we observed an increase in the % of Id4^+^/Ki67^-^ cells in Sox5^null^ in comparison with control NSCs in both conditions (FGF2: 5.7± 0.7 % vs. 2.8 ± 0.4 %; *P*= 0.004; FGF2+BMP4: 46.1 ± 3.1 % vs. 35.5 ± 2.7 %; *P*= 0.02; Fig. 7H, I). These results stress the point that Sox5 prevents an excess of pNSCs as observed in Fig. 7F. Finally, we found an intermediate population of Id4^+^/Ki67^+^ NSCs, which was increased in Sox5^null^ in the proliferating condition with respect to Control mice (7.2 ± 0.6 % vs. 3.3 ± 0.6 %; *P*= 0.0004; Fig. 7H, I), similarly to what we observed *in vivo* in Sox5^null^ mice at P14 (Fig. 3C-E). This population *in vitro* could represent the fraction of NSCs transitioning between the primed (Id4^+^/Ki67^-^ pNSCs) and the active states (Id4^-^/Ki67^+^ aNSCs; Fig. 7J). In summary, these results indicate that loss of Sox5 provokes by P14 an increase in pNSCs, a population that can be identified *in vitro* by cell tracer retention and high levels of BMP canonical signalling (including P-Smad1/5/9 and Id4 expression; Fig. 7J). These data also suggest that maintaining restricted levels of BMP canonical signaling during postnatal DG development could be essential to establish the correct number of pNSCs in the adult neurogenic niche that ensures long-lasting maintenance of the NSC pool.

## Discussion

To understand the process of adult neurogenesis is essential to determine how adult quiescent NSCs emerge during development and how they acquire a reversible state of quiescence. In this study, we have shown that Sox5 transcription factor: i) restricts NSC quiescence entry during the first postnatal week; ii) limits the establishment of a primed quiescent state during the second postnatal week and iii) restricts the levels of BMP canonical signaling pathway during DG postnatal development. Moreover, we have established that an excess of pNSCs leads to a hyper-neurogenic adult NSC pool that is prematurely exhausted. Thus, establishing the correct balance between dormant and primed quiescent sub-states during postnatal NSC development is essential for long-term maintenance of the neurogenic niche (Summary in Fig. 8).

**Fig. 8.**
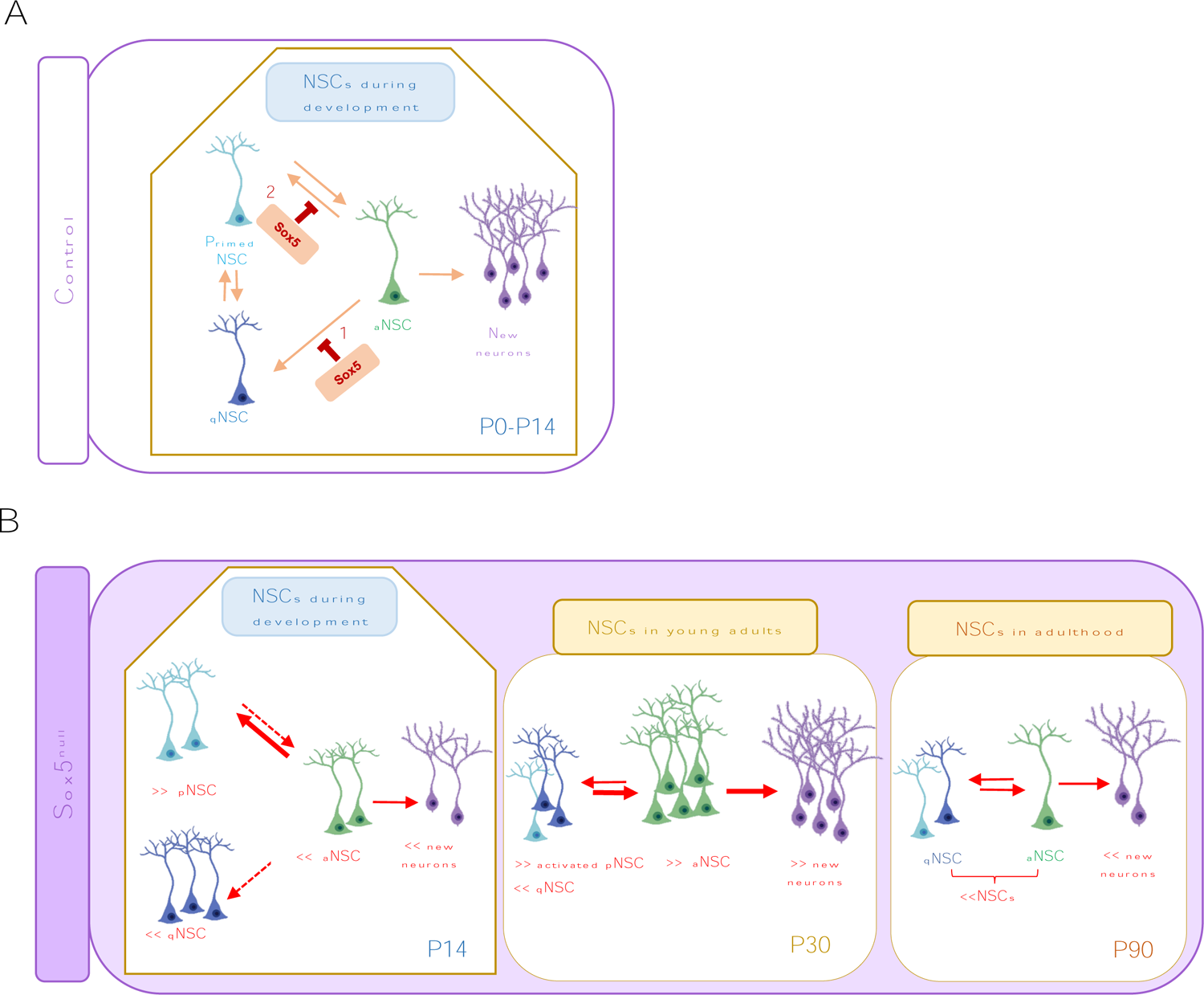
Summary of NSC population dynamics during early postnatal DG development and their alterations in Sox5^null^ mice. **(A)** During the first postnatal week NSCs enter into the quiescence state. We have shown that Sox5 transcription factor restricts NSCs quiescence entry during the first postnatal week (1). Moreover, around P14 NSCs acquire a superficial or primed state of quiescence and Sox5 limits this primed quiescence state (2). **(B)** The consequences of Sox5 loss during DG development are an increase in pNSCs and an initial reduction in both aNSCs and new neurons at P14. This is followed by an increase in the activation of the pNSCs and an excess in aNSCs that generate an overproduction of immature new neurons at P30. All these changes in young mice provoke a reduction in the NSC pool and a decrease in neurogenesis by P90

The first postnatal week of DG development is critical for the generation of adult NSCs. First, specific elimination of dividing NSCs between P0-P7, but not during P14-P21, severely impairs the size of the adult NSC pool [36]. Moreover, label-retaining birth-dating experiments revealed that NSC first entry into quiescence occurs during the first postnatal week [1,3] and that Sufu embryonic deletion (which decreases Shh signalling) promote premature quiescence entry during that week [3]. Using long-term retention of thymidine analogues, we now show that Sox5 is a new player in that early process, as it is required during the first postnatal week to prevent an excessive entry of NSCs into quiescence and to promote NSC proliferation. However, whereas Shh reduction leads to a maintained reduction in neurogenesis along time, we have observed that Sox5 loss has revealed another important quiescence modulation during the second postnatal week, when Sox5 is required to prevent an excess of NSCs in a shallow quiescent state as we describe below.

There is an emerging notion in the field that quiescent adult NSCs are at different levels of quiescence depth, forming a continuum of sub-states along the quiescence-activation trajectory. Single-cell and bulk transcriptomic analysis from adult SVZ and SGZ have revealed adult NSCs in intermediate states which includes a shallower or primed state [37,11,9,12,8], cell cycle-low activated state [38] or “resting” state [10]. However, it is unclear when do those intermediate states emerge during the neurogenic niche development. Our work indicates that NSCs acquired a primed state along the second postnatal week. Precisely, upon Sox5 loss, both NSC activation and dormant quiescence decreased around P14, whereas the number of NSCs at a primed state (pNSCS) is increased. These pNSCs *in vivo* do not retain BrdU-L for weeks (they are not dormant qNSCs) and co-express quiescence marker Id4 and proliferating marker ki67. We have reinforced this characterization by *in vitro* DFFDA cell tracer retention assays [11] showing that the pool of pNSCs in P14 Sox5^null^ mice is enlarged and is enriched in NSCs with activated BMP canonical pathway which includes expression of P-Smad1/5/9 and Id4 proteins. Consequently, we propose that around P14 there is a critical window to establish the correct balance between different sub-states of quiescence in the hippocampal NSCs and that it is essential to restrict the number of pNSCs. Moreover, as NSCs can only undergo a limited number of rounds of cell division prior to terminal differentiation [39], that early decision of preventing an excess of NSCs in a primed state is fundamental to establish long-lasting NSC pool and neurogenic capacity in adult mice.

Our results are reinforced by recent studies indicating that autophagy is required for the acquisition and maintenance of NSC quiescence in the first postnatal weeks [7] and by CcnD2 mutant analysis, which suggest that P14 NSCs are in a transition from developmental to adult NSC state [40]. The absence of cyclin D2 does not affect normal development of the DG until birth but prevents postnatal formation of adult NSCs which are born on-site from precursors located in the DG shortly after birth. Our analysis would indicate that the developmental to adult NSC transition could include the establishment of a primed state controlled in part by Sox5 activity.

Recent studies have described that the probability of adult NSCs to transit to a shallow or resting quiescence state increases over time, comparing one-month old and ageing DGs [10]. Ascl1 plays a relevant role in the maintenance of that resting state in adult NSCs as in *Ascl1* hypomorph mice there is an increase in the proportion of adult NSCs that return to quiescence after cell division and consequently an enlarged NSC pool [4,10]. Opposite to what has been described in Ascl1 deficient mice, in young adult Sox5^null^ mice pNSCs return less to quiescence after cell division and NSC activation increases. This excess of pNSC activation could be sustained by deep transcriptomic changes, including upregulation of cell cycle activators and MAPK pathway, as we have described. Moreover, those alterations cause a progressive reduction of the NSC pool and have a deleterious effect in elder Sox5^null^ mice neurogenesis. While Sox5 and Ascl1 may exhibit contrasting roles in regulating the return of pNSCs to quiescence in young adult mice, it remains unclear whether Ascl1 participates in the initial establishment of a primed state in developmental NSCs, akin to the role described here for Sox5.

Regarding possible mechanisms controlling the primed state, we have found that BMP canonical signalling pathway through P-Smad1/5/9 and Id4 expression could play a fundamental role in promoting the primed state in NSCs during the first postnatal weeks. BMP canonical pathway controls several aspects of DG development and function. Thus, during development inhibition of embryonic BMP canonical signalling in NSCs impairs DG neurogenesis [41]. Furthermore, BMP canonical pathway components such as BMP4 and its downstream effectors Id1 and Id4 are essential for quiescence maintenance in adult NSCs [13–17]. Now, we have demonstrated that Id4 expression is indicative of a primed state in NSCs when combined with Ki67 expression, both *in vitro* and *in vivo*. Moreover, Sox5 is crucial to prevent an excess of BMP canonical pathway activation in NSCs as indicated by our in vivo bulk RNA-Seq, RT-qPCR analysis and *in vitro* NSC immunostaining (Id4 and P-Smad1/5/9).

Our work also points to the fact that developmental loss of Sox5 could have severe consequences on hippocampal function, as new DG neurons promote flexible learning and adaptive behavioural and endocrine responses to cognitive and emotional challenges. Moreover, given the fact that *SOX5* heterozygous inactivating variants cause neurodevelopmental disorders in humans (Lamb-Shaffer Syndrome; OMIM # 616803), our findings will also help to understand the episodic and social memory deficits described in those patients [42].

## Materials and Methods

### Mice

All experiments were performed in postnatal three day-old (P0) to 5-months-old C57BL/6J background mice of both genders. Animal procedures were carried out in accordance with the guidelines of European Union (2010/63/UE), Spanish legislation (53/2013, BOE no. 1337) and under the approved projects (PROEX 078/17 and PROEX 146.0/22). Mice were housed with a standard control of a 12 h light/dark cycle and maintained in the animal facility at Cajal Institute.

Sox5^fl/fl^ mice, in which coding exon 5 of Sox5 gene is flanked by loxP sites were used [26]. For conditional mice, they were bred with Nestin-Cre mice (RRID:IMSR_JAX:003771) to generate control animals (Control): Nestin-Cre/Sox5^fl/+,^ Sox5^fl/fl^ or Sox5^fl/+^ and mutant mice: Nestin-Cre/Sox5^fl/fl^ (Sox5^null^) For conditional inducible mice, they were bred with Sox2-creER^T2^ mice [43] and with Rosa26-loxP-stop-LoxP-TdTomato (RRID: IMSR_JAX:007908) reporter mice to generate control animals (Control): Sox2-creER^T2^/TdTom^+^ or Sox2-creER^T2^/Sox5^fl/+^/TdTom^+^ and mutant mice: Sox2-creER^T2^/Sox5^fl/fl/^TdTom^+^ (Sox5^icKO^).

### Tamoxifen treatment and administration of Brdu/EdU

For activation of the CreER^T2^ protein, 5 mg/40gr body weight of tamoxifen (TAM, 10 mg/ml in corn oil; Sigma) was intraperitoneally (i.p.) injected once (in P3 mice) or three times, one injection every 24 h (in P14 mice). The labelling of cells progressing though S-phase was performed by either i.p. injections of Brdu (Roche) or EdU (Sigma Aldrich) (50 mg/kg) using PBS as vehicle or through administration of EdU in the drinking water (0.2 mg/ml). Animals were injected once (P3 and P14 mice) or five times with one daily injection (one month-old mice) and sacrificed 15 or 30 days after. For long-term (14 days) EdU administration in water, fresh solution was replaced at least every 56 hours.

### Tissue Preparation and Immunofluorescence

Animals were transcardially perfused with saline followed by 4% paraformaldehyde (PFA). Brains were postfixed with 4% PFA for 3 h at 4°C, embedded in 30% agarose/sucrose (w/v) and coronally sectioned in 50μm slices using a vibratome.

For immunostaining, vibratome floating brain sections were permeabilized with 1% Triton X-100 in 0.1M Phosphate Buffer (PB) for 30 min and blocked with 10% Fetal Bovine Serum (FBS) and 0.25% Triton X-100 in 0.1M PB for 2 h at room temperature with rocking. For fixed cells, permeabilization and blocking were performed with 10% FBS in 0.25% Triton X-100 in 0.1M PB for 1 hour at room temperature. Primary antibodies: BrdU (1/500, ab6326 Abcam), DCX (1/500, ab18723 Abcam), GFAP (1/1000, Z0334 Dako), Hopx (1/500, sc-398703 Biogen), P-Smad1/5/9 (1/500, ab92698 Abcam), P-Smad2^S467^(1/500, ab280888 Abcam) Id4 (1/500, BCH-9/#82-12 BioCheck), Ki67 (1/500, 550609 Bd Biosciences), MCM2 (1/1000, 610701 Bd Biosciences), Prox1 (1/1000, MAB5654, Millipore), Sox5 (1/500, A. Morales), Sox2 (1/500, AF2018 R&D), Tbr2 (1/1000, ab216870 Abcam), were prepared in incubation buffer (1% FBS, 0.25% Triton X-100 in 0.1M PB) and incubated with sections or cells overnight at 4℃. Following 3 washes, with /washing buffer (0.1% Triton X-100 in 0.1M PB), immunoreactivity was detected appropriate Alexa Fluor-conjugated (Life Technologies, Invitrogen and Abcam) secondary antibody (1:1000) diluted in incubation buffer for 2 h at room temperature. After three washes, sections or cells were incubated with bisbenzimide (1:100 in 0.1M PB) for 2 min at room temperature and mounted using Fluoromount-G (Thermo Fisher).

To detect MCM2 protein, antigen retrieval was carried out by incubating sections in 0.15M sodium citrate at 80 ℃ for 30min. To detect BrdU incorporation, DNA was denatured by incubating sections or cells with 2N HCL for 25min at room temperature, followed by 0.15M boric acid neutralization for 20 min. To detect Edu signal was used “EdU IV Imaging Kit 647 S” based on the Click-it reaction with azides conjugated to fluorochromes (Sigma Aldrich, BCK647-IV-IM-S) following instructions of the manufacturer.

### RNA extraction, library preparation and sequencing

DGs of Control and Sox5^null^ (n=4 and n=3 independent animals respectively) were dissected and RNA was extracted with the QuickGene RNA tissue kit S (Kurabo) following instructions of the manufacturer. Stranded mRNA libraries (enriched in polyA) were prepared, and the samples were sequenced using the platform HiSeq 2500 (Illumina) to an average output of 30 million 50bp single end reads per sample at CRG (Center of Genomic Regulation). Original data include 14 reads sets in FASTQ format. The seven samples were sequenced twice to ensure a number of reads greater to 30 million.

### RNA-Seq and differential expression analysis

Next-generation sequencing (NGS) data bioinformatic analysis has been performed by the Genomics and NGS Core Facility (GENGS) at CBMSO (CSIC-UAM) Madrid, Spain. Quality analyses were performed over reads using FastQC1 (v0.11.8) software. The quality by position (Phred+33 quality score) maintained in general good standard across all cycles with median and mean base quality over 28. Reference genome and the annotation file of *Mus musculus* (GRCm39, mm39) have been downloaded from UCSC ftp site and GENECODE respectively. The reads were aligned against *M. musculus* genome using Hisat24 (v2.1.0) aligner. Before the alignment, the FASTQ files from the same sample, but generated each from different sequencing runs, were merged. Results showed a good behaviour of reads in the alignment process, on average more than 96 % of reads mapped against the reference genome.

Integrative Genomics Viewer [44] was used to visualize aligned reads and normalized coverage tracks [RPM (reads per million)]. We used htseq-count7 (v0.11.2) to count the reads mapping each feature. We have used the “intersection-strict” resolution mode, where reads are counted only if they are inside a gene or inside the exons of a gene. The differential expression analysis was performed using Deseq2, an R software package, and differentially expressed genes (DEG, with adjusted p-value < 0.05) were represented in heat map graph and volcano plot. Specific heat maps of signalling pathways of interest were also generated using the same Bioconductor package. RNA-Seq data reported in this study are accessible through the ENA accession number PRJEB48583 0e3278a4-42af-4e40-acb3-cda6b8350562.

### Gene set enrichment and pathway analyses

Identification of enriched biological functions and processes in DEGs was performed using gene enrichment analysis and gene ontology (GO) analysis using freely accessible Panther software (version 17.0). The PANTHER classification system is a comprehensive system that combines genetic function, ontology, signalling pathways, and statistical analysis tools to study large-scale experimental genome-wide data and its relationship to certain previously annotated processes. We analysed GO terms for cellular processes, cellular components, and molecular functions, as well as overrepresented signalling pathways in our DEGs. We used the PANEV software (Pathway Network Visualizer) [33], which consists of a set of R packages, for the visualization of gene set/pathway-based networks. This software is based on information available in the KEGG database and visualizes genes within a network of interconnected upstream and downstream pathways at multiple levels (from level 1 to n). Network visualization helps interpret the functional profiles of a group of genes. We used this software to construct networks of cell cycle and TGF-β/BMP signalling pathways.

### RNA extraction, cDNA synthesis and real-time quantitative PCR

Total RNA of Control and Sox5null DG at P34 were extracted using the QuickGene RNA tissue kit S (Kurabo) and then treated with DNAse. cDNA was synthesized with SuperScriptTM IV First-Stand Synthesis System and random hexamers (Invitrogen). Gene expression levels were measured using TaqMan Gene expression assays (Applied Biosystems) and quantitative real-time PCR (RT-qPCR) was carried out in a 7500 PCR System using TaqMan Fast Advanced Master Mix (Applied Biosystems). The following Taqman assays probes were used: ACVR1 (Mm01331069_m1), BMP3 (Mm00557790_m1), Bmpr1a (Mm00477650_m1), Bmpr1b (Mm03023971_m1), Egr (Mm00456650_m1), Fos (Mm00487425_m1), Gapdh (Mm99999915_g1), Id2 (Mm00711781_m1), Id4 (Mm00499701_m1), Sox5 (Mm01264584_m1) and SMAD7 (Mm00484742_m1). Gene expression was measured relative to endogenous control Gapdh and normalized to the expression of the Control sample in each group using the 2-ΔΔCt method, as indicated in the corresponding figure. At least three independent experiments were performed for each condition and samples were run in triplicates.

### NSCs culture and in vitro treatments

Postnatal P14 hippocampal NSCs were cultured as previously described [24]. Briefly, mice were euthanized with CO2, their brains were isolated, and the hippocampus were dissected, cut up into pieces and digested with Papain [0.66 mg/ml papain (Worthington) + 0.2mg/ml cysteine (Sigma) + 0.2 mg/ml EDTA (Merck) + Hank’s buffer (Thermo Fisher)] for 15 min at 37°C. After mechanical dissociation and washes with DMEM F12 (Thermo Fisher) to stop the reaction and washes with Hank’s, the disaggregated cell suspension was plated into MW12 plates with basal media [DMEM F12 + 1X N2 supplement (100X; Thermo Fisher) + 1X B27 supplement (50X; Thermo Fisher)], 20 ng/ml EGF (100 ng/ μl; PeproTech) and 20 ng/ml FGF2 (100 ng/ μl; PeproTech). Cells were incubated at 37℃ and 5% CO2. Normally, one single brain was used to prepare the culture and 4 wells of MW12 per brain were used. For hippocampal floating neurospheres, 20 ng/ml EGF and 20ng/ml FGF2 were daily added and were passaged by mechanical procedures and used from passage 0 until passage 15 for different cell treatments. For immunostaining experiments in intact neurospheres we used Matrigel coated glasses to attached grown neurospheres during 10-15 min at 37 °C. To prepare matrigel coated glass, clean glass coverslips, were coated with diluted Matrigel (Thermo Fisher) in DMEM (1/100) in MW24 and incubated 4-12 h at 37°C.

For the analysis of fluorophores retention, neurospheres were dissociated into a single cell suspension and incubated in 0.5 mL of PBS with 2 mg/ml Cell Trace Oregon Green 488 Carboxy-DFFDA SE (Thermo Fisher) for 7 minutes at 37°C in the dark as previously described [11]. After that, cells were washed with control medium, centrifuged at 200g for 10 min, resuspended and seeded in NSC complete medium. After 6 days, neurospheres were dissociated and the fluorescence intensity of cell tracer was measured in a flow cytometer (BD).

### Microscopic Analysis and Cell counting

All images were taken with a direct SP5 confocal microscope (Leica). Images of both left and right dorsal DG sections (−0.82 mm to −4.16 mm from bregma) were captured with a z-step of 2 μm through at least 20 μm of each 50 μm sections. Labelled cells were counted in the SGZ of every ninth of 50 µm DG sections. In Control and Sox5^null^ mice the analysis was done counting the number of cells (at least 300 cells/marker) that expressed a cell-type specific maker in the population of cells among SGZ cells expressing a certain cell-type marker. In Control and Sox5^icKO^ mutant mice we counted recombined TdTom^+^ cells that were positive for the indicated marker. In those cases, 3 to 6 sections from at least 3 to 7 mice and a minimum of 100 cells for each animal were analyzed. Counting was performed manually and blind using LAS X (Leica) software. In all the cultures, three independent experiments were performed for each condition.

### Statistical Analysis

The appropriate sample size (N) was determined based on similar published data from other groups [10], using a minimum of 3 mice per condition for in vivo experiments, and a minimum of biological triplicate for in vitro experiments. Statistical analysis and graphs were conducted with GraphPad Prism version 8 software using different tests with a significance level of at least p< 0.05. Thus, two-tailed unpaired Student’s t test were used for most of statistical comparisons of two conditions for in vivo experiments; paired t test for in vitro experiments where control and treatment conditions for each biological replicate were performed in parallel. Statistical details were included in each figure and figure legend [number of experiments (N), number of cells (n) and statistical test]. Data are presented as mean ± SEM. Significance is stated as follows: p<0.05 (*), p<0.01 (**), p<0.001 (***), confidence intervals of 95%.

## Supporting Information

Figures in a separated Pdf file.

**S1 Fig. Sox5 is expressed in Sox2 expressing progenitors and NSCs during DG development.** (**A,B**) Confocal images showing Sox5 and Sox2 immunostaining in E16.5 (A) and P0 (B) mouse hippocampal sections. CA, Cornus Ammonis; CH, cortical hem; DG, dentate gyrus. (C) Confocal images showing Sox5+ cells in dorsal DG in P5 and P14 mice and in combination with the indicated markers in P5 mice. (D, E) Quantitation of positive cells for the indicated marker amongst Sox5+ (D) or Sox2+ population (E) in P5 and P14 mice. At least 3 animals and 3 sections/animal were analyzed for each immunostaining. In all graphs, data are mean value ± SEM. ***p < 0.001 by unpaired Student’s t test. Scale bar represents 100 µm (A left, B right), 150 µm (B left, C left) and 25 µm (A right, F right)

**S2 Fig. Loss of early postnatal Sox5 expression causes an increase in the activation of dormant qNSCs in young adult mice. (A)** Experimental scheme of EdU label retention experiment in P5>P30 in Sox5^icKO^ mice to label all pNSCs and aNSCs (EdU+) during the P30>P45 period. EdU^-^ NSCs could correspond to dormant qNSCs and EdU^+^ NSCs to pNSCs and recent aNSCs. (**B**) Confocal images showing TdTom expression and Nestin, Ki67 and EdU immunostaining in the SGZ of P30 Control mice after the procedure described in (A). (**C**) Quantitation of % of different NSC subpopulations (recombined Nestin^+^Tdtom^+^) including pNSCs that have returned to quiescence (EdU^+^Ki67^-^) or that have been activated (EdU^+^Ki67^+^), deep dormant (EdU^-^Ki67^-^) and activated dormant qNSCs (EdU^-^ Ki67^+^) in the EdU+ or EdU-NSC population of P45 Control (n= 5) and Sox5^icKO^ (n=3) mice. In the graph, data represent mean values ± SEM. *p < 0.05 by unpaired Student’s t test. Scale bar represent 10 µm in B

**S3 Fig. Loss of Sox5 expression causes an early reduction in neurogenesis by P14.** (A) Confocal images showing DCX and BrdU immunostaining in the SGZ of DG at the indicated stages in Control and Sox5^null^ mice. (B) Scheme of BrdU labelling of newborn neurons from P12 to P14. (C) Quantitation of DCX+ BrdU^+^ cells/mm^2^ in P14 Control and Sox5^null^ mice. In the graph, data are mean value ± SEM. *p < 0.05 by unpaired Student’s t test. Scale bar represent 25 µm in A

**S4 Fig. Analysis of neurosphere number and diameter in P14 Control and Sox5^null^ mice DG in acute NSC preparation. (A)** Acute NSCs preparation from P14 DG, from passage 0 (primary, 1^ry^) to passage 5. Quantitation of neurosphere number per area for 1^ry^ and 2^ry^cultures, and in successive passages for Control and Sox5^null^ mice. **(B)** Quantitation of neurosphere diameter in 1^ry^, 2^ry^ and 3^ry^ cultures for Control and Sox5^null^ mice. Data are mean value ± SEM. ***p < 0.001 by unpaired Student’s t test

## Acknowledgments

We thank Inés Colmena and Carmen Capitán for her technical assistance.

## Author Contributions

AVM conceived the project and together with LL and CMM designed the experiments, interpreted the data and wrote the manuscript. LL, CMM, PRM, PTM, EMP, RLS and MVB performed experiments and analyzed the data; AVM, LL and CMM supervised experiments and discussed results. All the authors contributed to and approved the manuscript.

## Data availability

Microscopy, cell counting, RT-qPCR, and cell transduction data reported in this paper will be shared by the lead contact upon request. RNA-Seq data reported in this study are accessible through the ENA accession number PRJEB48583 0e3278a4-42af-4e40-acb3-cda6b8350562. Any additional information required to reanalyze the data reported in this paper is available from the lead contact upon request.

## Abbreviations

aNSC: active neural stem cell

BrdU-L: BrdU-long retaining NSCs

DEG: differentially expressed genes

DG: dentate gyrus

GN: granule neuron

IPC: intermediate progenitor cell

NSC: neural stem cells

pNSC: primed neural stem cell

qNSC: quiescent neural stem cell

RT-qPCR: quantitative reverse transcription polymerase chain reaction

SGZ: subgranular zone

TAM: tamoxifen.

